# LPA/PKD-1 signaling promotes development of arteriolar niche that supports self-renewal of breast cancer stem cells and stemness

**DOI:** 10.1101/2020.10.22.351122

**Authors:** Yinan Jiang, Yichen Guo, Jinjin Hao, Rachael Guenter, Justin Lathia, Adam Beck, Reagan Hattaway, Doug Hurst, Q Jane Wang, Yehe Liu, Qi Cao, Helen Krontiras, Herbert Chen, Roy Silverstein, Bin Ren

**Author notes:** **Correspondence to** Bin Ren, MD, PhD, FAHA, Department of Surgery & O’ Neal Comprehensive Cancer Center, University of Alabama at Birmingham School of Medicine. 1824 6^TH^ Ave South, Wallace Tumor Institute WTI630C2 (O) & WTI620E (L), Birmingham, AL35294. These authors contribute equally to this work (Yinan Jiang, Yichen Guo, Jinjin Hao).

## Abstract

Breast cancer stem cells (BCSCs) are essential for cancer growth, metastasis and recurrence. However, the regulatory mechanisms of self-renewal and interactions with the vascular niche within tumor microenvironment are currently under investigation. Here, we demonstrate that BCSCs are enriched within arteriolar niche within the tumor microenvironment of estrogen receptor positive (ER^+^) BC and bi-directionally interact with arteriolar endothelial cells (ECs). Mechanistically, this interaction is driven by the LPA/PKD-1 signaling pathway, which promotes arteriolar differentiation and self-renewal. Furthermore, this pathway directly promotes stemness features. These findings suggest that targeting LPA/PKD-1 signaling may disrupt the arteriolar niche within the tumor microenvironment and concomitantly eradicate BCSCs, thereby attenuating BC progression.

## INTRODUCTION

In many advanced cancers, recurrence and metastasis is driven by a subpopulation of cells called cancer stem-like cells (CSCs). These cells are characterized by their enhanced tumor-initiating and self-renewal capacities (1). Al-Hajj and colleagues initially identified tumorigenic CSCs in breast cancer (BC) known as BCSCs (2). The maintenance and expansion of BCSCs depends on both cell intrinsic pathways and the interaction with the surrounding tumor microenvironment. Tumors harbor CSCs in dedicated niches (3–8), including vascular niches in BCs (9, 10). Niche stromal cells, such as tumor-associated endothelial cells (TECs), serve multiple functions within the tumor microenvironment (11). Effective inhibition of angiogenesis in the tumor microenvironment, a hallmark of BC progression (12), could disrupt the vascular niche and render BC dormant (13). Unfortunately, current anti-angiogenic therapies are ineffective under experimental conditions (14–16) and show limited efficacy in the clinic (17–19). The resistance to anti-angiogenic therapy may result from resistance to an established mature vasculature (16, 20), indicating that the current strategies targeting the vascular niche should consider heterogeneity in tumor vasculature (20–22).

Aberrant tumor vascularity has repeatedly been shown to be important for satisfying voracious demands for nutrients by rapidly growing tumors, permitting metastasis, and serving as abnormal CSC niches (23, 24). Arterial networks may exist in solid tumors (21, 22, 25) to provide oxygen and nutrients for tumor progression (19, 22, 26, 27). Furthermore, studies have shown that liver and lung metastases are arterialized (28, 29). However, it remains largely unknown if there is an arteriolar niche in the tumor microenvironment of BCs and how heterogeneous blood vessels and perivascular niches participate in the regulation of BCSCs. Furthermore, the signaling mechanisms that regulate the arteriolar niche and the stemness of BCSCs are also largely unclear.

There have been multiple signaling pathways essential for angiogenesis and pro-arteriogenic reprogramming of ECs, including lysophosphatidic acid (LPA)/protein kinase D (PKD-1), that likely regulate the vascular niche within the tumor microenvironment, in addition to promoting BC progression (30–34). LPA as a lipid signaling mediator may promote arteriolar differentiation by upregulation of such arteriogenesis-associated genes as ephrin B2 and DLL4 (35) via PKD-1 pathway (36). Recent studies suggest that this pathway also regulates the stemlike properties of cancer cells including BC cells (37–39). It is well-known that CD44 is a critical marker in BCSCs (2) whereas ALDH1 is also a CSC marker that promotes BCSC plasticity, metastasis, and progression (2, 40–44). Additionally, CD36 is considered not only an angiogenesis regulator in ECs (30, 34, 45) but also a driver for tumor progression and metastasis-initiating cell in CSCs (46, 47).

Based on these observations, we hypothesized that the LPA/PKD-1 signaling axis regulates both arteriolar differentiation of the vascular niche in the tumor microenvironment and self-renewal of BCSCs. In this study, we observed that BCSCs are enriched in the arteriolar niche of BC, a location where bi-directional interactions occur between arteriolar ECs and BCSCs. Moreover, we found that the LPA/PKD-1 signaling axis promotes arteriolar differentiation and self-renewal of BCSCs, two aspects that may be essential for BC progression and therapeutic relapse.

## RESULTS

### Enrichment of breast cancer stem-like cells in the arteriolar niche

To determine the distribution of BCSCs in the tumor microenvironment, we interrogated both the presence and localization of these cells in BC tissues from patients using immunohistochemistry. BC cells that demonstrate a marker profile of CD44^+^7CD24^-^7ALDH1A^+^ are commonly accepted as BCSCs (2). We detected CD44^+^/CD24^-^7ALDH1A1^+^ BCSCs within the tumor microenvironment (Supplementary Figure 1).

To more precisely define the location within the arteriolar niche where BCSCs tend to be enriched, we stained BC tissues from patients for the presence of ALDH1A1, ephrin B2 (an arterial EC marker essential for arterial function), and α-smooth muscle actin (α-SMA) for smooth muscle cells in arterioles (Figure1 A-D). ALDH1A1 served as a marker for BCSCs, whereas the costaining of ephrin B2 and α-SMA served as markers of the arteriolar niche. Intriguingly, there were significantly more ALDH1A1^+^-BCSCs localized to the arteriolar niche than in the tumor nest (*p* < 0.01; Figure 1D). In a preclinical setting, syngeneic mice serving as models of BC were treated with LPA and found to have expanded arteriolar networks as supported by a significant increase in α-SMA^+^ blood vessels (Supplementary Figure 2). Our current data suggest that ALDH^+^ CSCs that are localized within the arteriolar niche may show plastic (42, 44), which may be important in BC progression.

**Figure 1.**
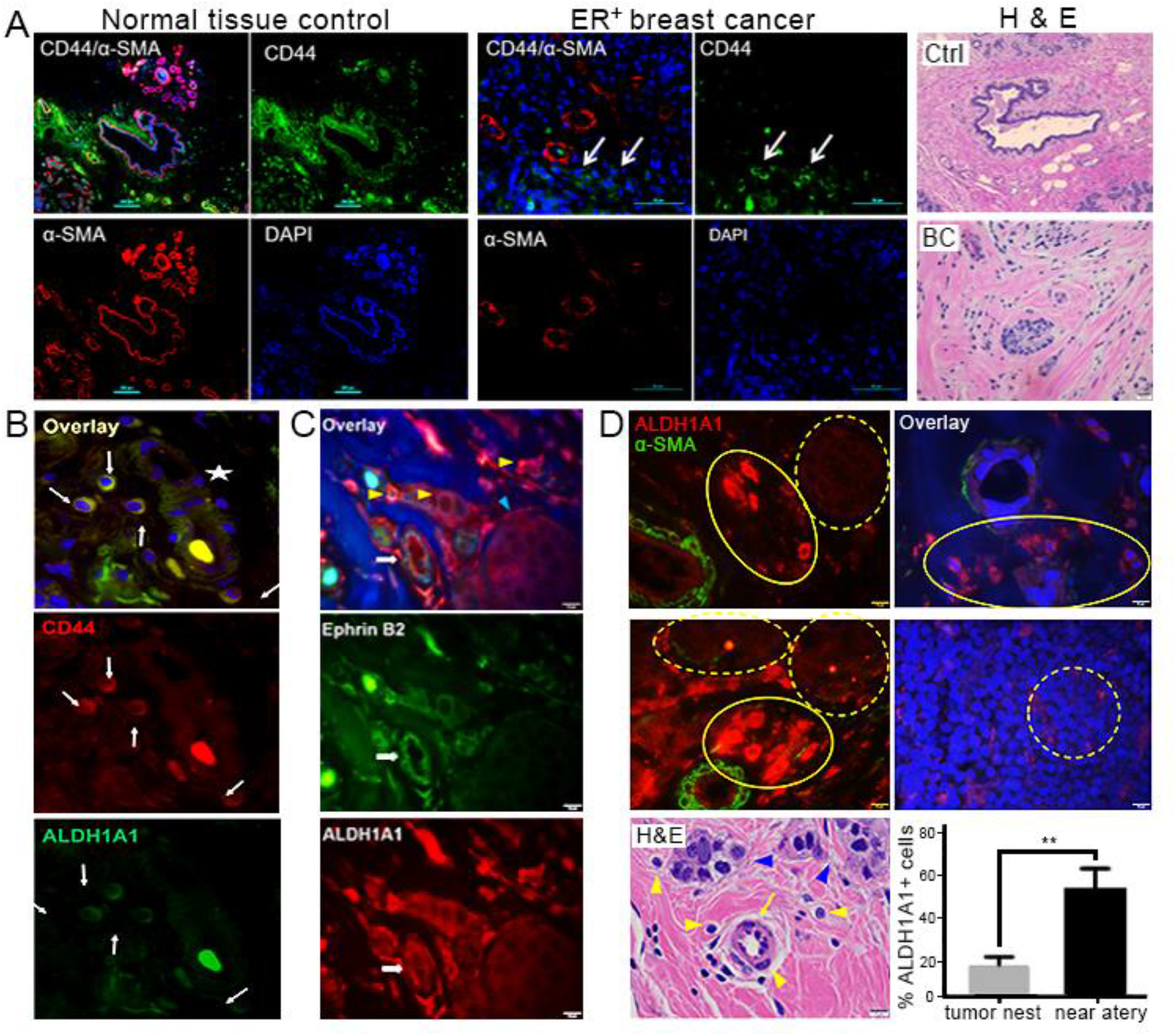
Breast cancer stem-like cells are highly enriched within the arteriolar niche. **A**. CD44^+^ cancer cells within vascular niche (white arrows), as shown by staining with α-SMA (red) in human ER^+^ breast cancer. DAPI (blue) is used for nuclei staining. Tissue structures are shown by H & E staining. **B.** BCSCs in a human BC specimen within TME. Human ER^+^ BC specimen was co-stained with anti-CD44 and anti-ALDH1A1 antibodies, followed with appropriate 2^nd^ antibodies and mounted with DAPI. Images were acquired using an Eclipse T*i*2^®^Nikon microscope and overlaid using NIS Elements software. BCSCs are indicated by white arrows and the blood vessel-like structure is shown by a white star. No CD24 was detected. Shown are representative images. Bar = 10 μm. **C.** Ephrin B2 and ALDH1A1 costaining in human ER^+^BC specimens. Ephrin B2 was stained green and ALDH1A1 red. White arrows show ephrin B2 positive arteriole, yellow arrow heads indicate ALDH1A1^+^ cancer cells, and tumor nest is indicated by light blue arrowhead. Cells in the cancer nests did not show significant ALDH1A1 staining. Bar = 10 μm. **D**. Quantification of ALDH1A1^+^ BC cells within the arteriolar niche. The arteriole is circled by yellow circles. BC cells were randomly counted under Olympus BX60 fluorescence microscope linked with CCD camera. The standard for examining cell locations were: 1) closed to artery is defined as ALDH1A1^+^ BC cells within 50 μm from the SMA^+^ small arteries, and 2) close to the cancer nest refers to ALDH1A1^+^ BC cells locating cancer cell nest over 50 μm from the SMA^+^ small artery. Up to 30 cancer cells were randomly counted within 50 μm x 50 μm area with five repetitions. \*\**P* < 0.01.

### Crosstalk between arteriolar ECs and BCSCs

To further investigate the relationship between BC cells and the arteriolar niche, we sought to determine whether BC cells interact with surrounding arteriolar ECs indirectly, directly, or both. As a test for indirect crosstalk, we used transwell chambers to co-culture estrogen receptor positive (ER^+^) BC cells (MCF-7) and human microvascular ECs (HMVECs) that express Deltalike ligand 4 (DLL-4), a marker of well-differentiated arteriolar ECs and ligand involved with Notch1 signaling. ER^+^ BC cells grown in co-culture with DLL4^+^-HMVECs had higher levels of gene expression of cancer stemness-related genes *(CD44, ALDH1A1, KLF4,* and *CD36)* and *PKD-1*, as compared to ER^+^ BC cells grown in monoculture (Figure 2A, \**p* < 0.05, \*\**p* < 0.01, \*\*\**p* < 0.001). These results suggest that BCSCs may take advantage of the arteriolar niche for their maintenance and self-renewal.

**Figure 2.**
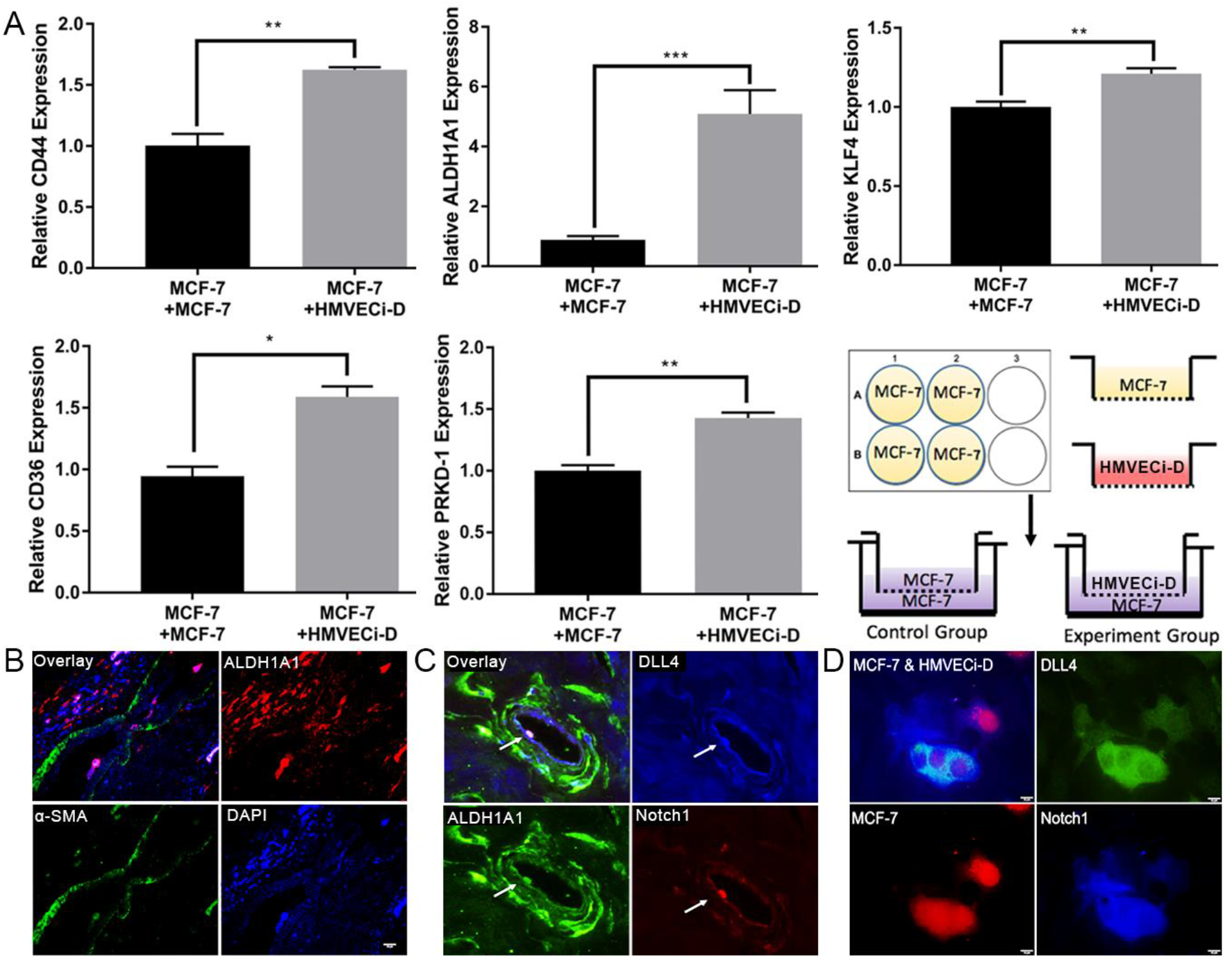
Indirect and direct interactions of breast cancer stem-like cells with the arteriolar endothelium within TME. **A.** Co-culture of breast cancer cells and endothelial cells increased expression of BCSC markers and stemness-associated genes. mRNA in MCF-7 cells cultured in 6-well plate was collected for detection of stemness-associated genes by RT-qPCR. The results of triplicate experiments are shown as mean ± SEM. \*\*\**P*<0.001, \*\**P*<0.01, or \**P*<0.05. **B.** ALDH1A1^+^ BCSC was attached to the arteriolar endothelium within the vascular lumen. Human BC specimen was stained with ALDH1A1 and alpha smooth muscle actin (α-SMA) antibodies used, followed by proper secondary antibodies and mounted with DAPI. Bar = 10 μm. **C.** ALDH1A1^+^ cell that expresses high levels of Notch1 and DLL4 interacts with DLL4^+^ endothelium in the arteriolar vessel. Human breast cancer specimen was processed and stained with Notch1, DLL4 and ALDH1A1 antibodies followed with appropriate secondary fluorescence antibodies. Bar = 20 μm. **D.** MCF-7 cells transduced with DsRed interacted with an HMVEC. MCF-7 cells transduced with Ds-Red that were grown in complete MammoCult™ Media for 5 days were cocultured with HMVECi-Ds that expressed with DLL4 at the ratio of 2:1 in DMEM for 48 hours. The cells were stained with Notch1 and DLL4 antibodies, followed with appropriate secondary fluorescence antibodies. Bar = 10 μm. The images were taken with an Olympus BX60 fluorescence microscope, and representative images are shown.

Additionally, PKD-1 signaling may play a dual role in both endothelial cells and BC cells to promote BCSC expansion (Figure 4D & E). To test for direct crosstalk, we three-dimensionally co-cultured (Supplementary Figure 3A) GFP-labeled BC cells (E0771) (Supplementary Figure 3B) with Ds-Red-labeled HMVECs (Supplementary Figure 3C) and observed the interactions of MVEC and BC cells in the margin area (Supplementary Figure 3A). Notch1 is an important molecule in human BC that regulates SC self-renewal (48, 49) and EC Notch1 promotes metastasis (50) in addition to its role in arteriolar differentiation (51, 52). We then characterized BCSCs present in the tumor vasculature *in vivo.* Using human ER^+^ BC specimens analyzed by immunofluorescence microscopy, we found the existence of Notch1^+^ BC cells within the microvasculature, suggesting a direct interaction with arteriolar ECs, which was supported by a vascular-like structure consisting of DLL4+ and DLL4+/Notch1+ cells (Supplementary Figure 4). Moreover, an ALDH1A1+ BC cell directly attached arteriolar endothelium (Figure 2B).

DLL-4, the ligand to the Notch1 receptor, has been shown to be upregulated in tumor vasculature (53), whereas this ligand is also expressed by arteriolar ECs in the vascular system (51,54). To confirm if the Notch1 pathway is involved in mediating the crosstalk between arteriolar endothelium and BCSCs, we analyzed BC tissue from patients for the expression of ALDH1A1, Notch1, and DLL-4. Interestingly, ALDH1A1+ BCSCs that co-expressed Notch1 showed direct contact with the tumor endothelium that expressed DLL4 in a tumor arteriole (Figure 2C). This interaction indicates a direct crosstalk between arteriolar ECs and BCSCs. As further confirmation of this interaction, we co-cultured enriched-BCSCs with MVECs that showed high levels of DLL-4 expression. The results demonstrated that Notch1^+^ BC cells were likely to directly interact with the DLL-4^+^ MVECs (Figure 2D). These data suggest that arteriolar differentiation may bridge the crosstalk between BCSCs and ECs via Notch1 signaling.

### LPA/PKD-1 signaling in arteriolar differentiation of vascular endothelial cells

Arteriogenic gene regulation and arterial differentiation are regulated by MAP kinase/Erk signaling ^(55–58)^, while Erk-related PKD-1 is integral to angiogenesis through its interactions with the VEGF and CD36 signaling pathway ^(34, 59–61)^. PKD-1 not only activates Erk(62) but also transduces PLCγ1 signaling^(63)^, also crucial in arterial differentiation^(64)^.

The lipid signaling mediator LPA activates PKD-1 to regulate angiogenesis by suppressing transcription of CD36, a well-established angiogenic regulator that initiates antiangiogenic responses ^(65)^ and mediates ischemic injury ^(66)^. To determine whether LPA/PKD-1 signaling is essential for arteriolar differentiation of vascular ECs, we treated HMVECi-D cells with either LPA and/or a PKD inhibitor and then examined the expression of arterial genes by RT-qPCR. Exposure to LPA significantly increased the gene expression of ephrin B2 compared to the control, an effect that was reversible upon treatment by a PKD inhibitor (Figure 3A, *\*p* < 0.05, \*\**p* < 0.01). To further define the role of PKD-1 signaling in the regulation of arterial gene expression, we isolated primary lung ECs from both wild type control (Flox) mice and the endothelial specific-*pkd-1* knockout mice. We found that the gene expression of PKD-1 was significantly downregulated in primary lung ECs from EC-specific *pkd-1* knockout mice (Figure 3B) whereas the expression level of CD36, an angiogenesis regulator in ECs that may participate in arteriolar differentiation (30), was significantly increased (Figure 3C). Correspondingly, the expression levels of such arteriogenic gene as ephrin B2, DLL-4, and neuropilin 1 were all significantly decreased compared to controls (Figure 3D). Western blotting further confirmed the decreased expression of ephrin B2 and neuropilin 1, along with the increased expression in CD36 in PKD-1-knockout lung ECs (Figure 3E). Furthermore, the protein expression of DLL-4 decreased, although it did not reach statistical significance (Figure 3E). These studies suggest that LPA/PKD-1 signaling is associated with arteriolar differentiation.

**Figure 3.**
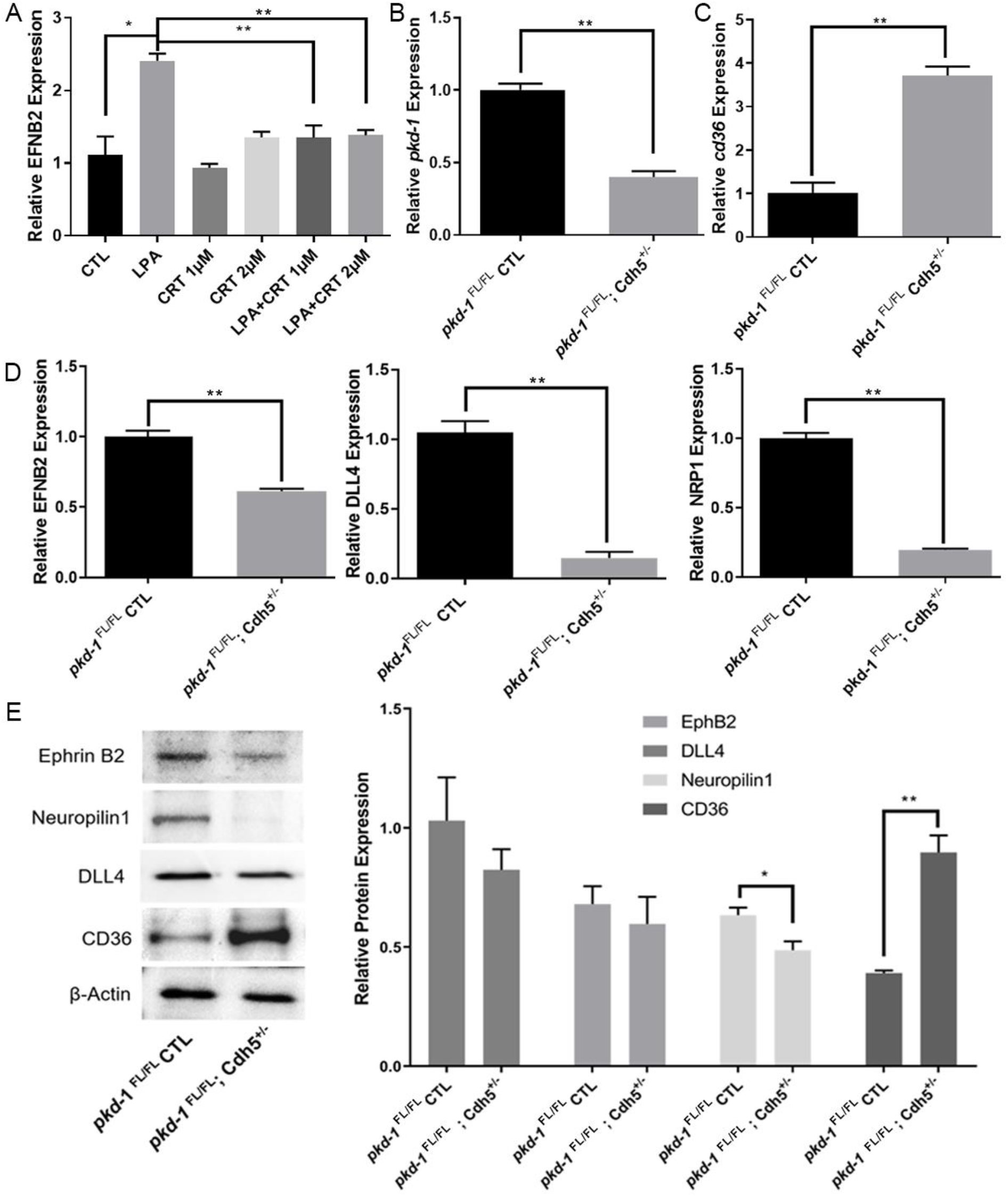
Regulation of CD36 and arteriogenic gene expression via PKD-1 signaling in vascular endothelial cells. **A.** HMVECi-D cells were cultured in endothelial medium (DMEM medium with 5% MVGS and 5% FBS). After starvation in serum free DMEM medium for 6 – 8 hours, the cells were treated with 10 μM of LPA, and/or 1 μM or 2 μM of CRT0066101 in serum free DMEM medium for 24 hours under 5% CO_2_ and 37°C. The total RNA was extracted and the mRNA levels of arterial gene ephrin B2 were detected by RT-qPCR. **B.** Primary lung endothelial cells (ECs) were isolated from the control and EC-specific *pkd-1* knockout mice. Passage 1 ECs were used for extraction and purification of total RNA. mRNA levels of *pkd-1* were detected by RT-qPCR and relative expression of *pkd-1* was compared between the control and *pkd-1-deficient* ECs. **C.** Increased expression of *cd36* in *pkd-1*-deficient lung ECs. Lung ECs were isolated from the control and EC-specific *pkd-1* knockout mice. Passage 1 lung ECs were used for extraction and purification of total RNA. mRNA levels were detected by qPCR and relative expression was compared between ECs from the control and EC-specific *pkd-1* knockout mice. **D.** Decreased expression of arteriogenic genes in *pkd-1*-deficient lung ECs. Primary lung ECs were isolated from the control and EC-specific *pkd-1* knockout mice. Passage 1 ECs were used for extraction and purification of total RNA. mRNA levels were detected by RT-qPCR and relative expression was compared between the control and *pkd-1*-deficient ECs. **E.** The cell lysate was extracted from passage 1 primary lung ECs that were isolated from EC-specific PKD-1 and control mice for detection of protein expression by Western blotting. Triplicate experiments were performed and levels of protein expression assessed by densitometry with Image J. * *P* < 0.05 or ** *P* < 0.01 *vs* control.

### LPA/PKD-1 signaling in self-renewal of BCSCs

In cancer compartment, LPA signaling may protect mesenchymal SCs from apoptosis(67) and promote the expansion of CSCs (37), which contributes to tumor progression (68), including ER^+^ BC (31, 69). To test the hypothesis that LPA signaling promotes the expansion of BCSCs via PKD-1, we examined the expression of PKD-1 in a syngeneic BC model with or without administration of LPA. Immunohistochemical assays demonstrated that LPA treatment significantly increased PKD-1 expression in BC tissues (Supplementary Figure 5). To determine the expression level of PKD-1 in patients, we examined pathological specimens resected from patients with ER^+^ BC. Intriguingly, PKD-1^+^ cancer cells presented a distinct distribution between different patients. A subpopulation of BC cells showed high levels of PKD-1 expression, particularly in the tumor nests (Supplementary Figure 6A), whereas singular BC cells had moderate expression of PKD-1 (Supplementary Figures 6A-D & 7).

To validate the association of PKD-1 signaling with BC stemness *in vivo,* we utilized immunofluorescence microscopy. As expected, we identified BC cells that were positive for both PKD-1 and CD44 in a syngeneic mouse BC model (Supplementary Figure 6B) and in BC patient’s specimens (Supplementary Figure 6C). Interestingly, the individual BC cells with positive costaining for both PKD-1 and CD44 were located outside of the tumor nest in patient BC specimens (Supplementary Figure 6C).

Furthermore, the Oncomine database analysis (70) demonstrated that the expression of CD36, a driver for tumor progression and metastasis (31, 46, 47, 71) was associated with tumorigenesis in an aggressive ER^+^ BC cell line (Supplementary Figure 8A). In this study, CD36 levels were increased in individual cells or tumor nests in ER^+^ BC patient specimens (Supplementary Figure 6D & Supplementary Figure 8B). Moreover, a few CD36^+^ cells existed in ALDH1A1^+^- and CD44^+^-mouse BC cells that were transduced with GFP (Supplementary Figure 6E). Similarly, a few BC cells co-stained positively for both PKD-1 and CD36 in mouse BC cells (Supplementary Figure 6F).

Intriguingly, a subset of CD44^+^ BC cells with high expression levels of PKD-1 and CD36 were localized within or nearby blood vessels and tumor nests (Figure 4A & B). BCSCs with high CD36 and moderate PKD-1 expression were localized outside of the tumor nest and had invaded into the surrounding tissue (Figure 4C), whereas the BCSCs with low CD36 and moderate PKD-1 expression appeared to form a vessel-like structure or capillary (Figure 4C).

**Figure 4.**
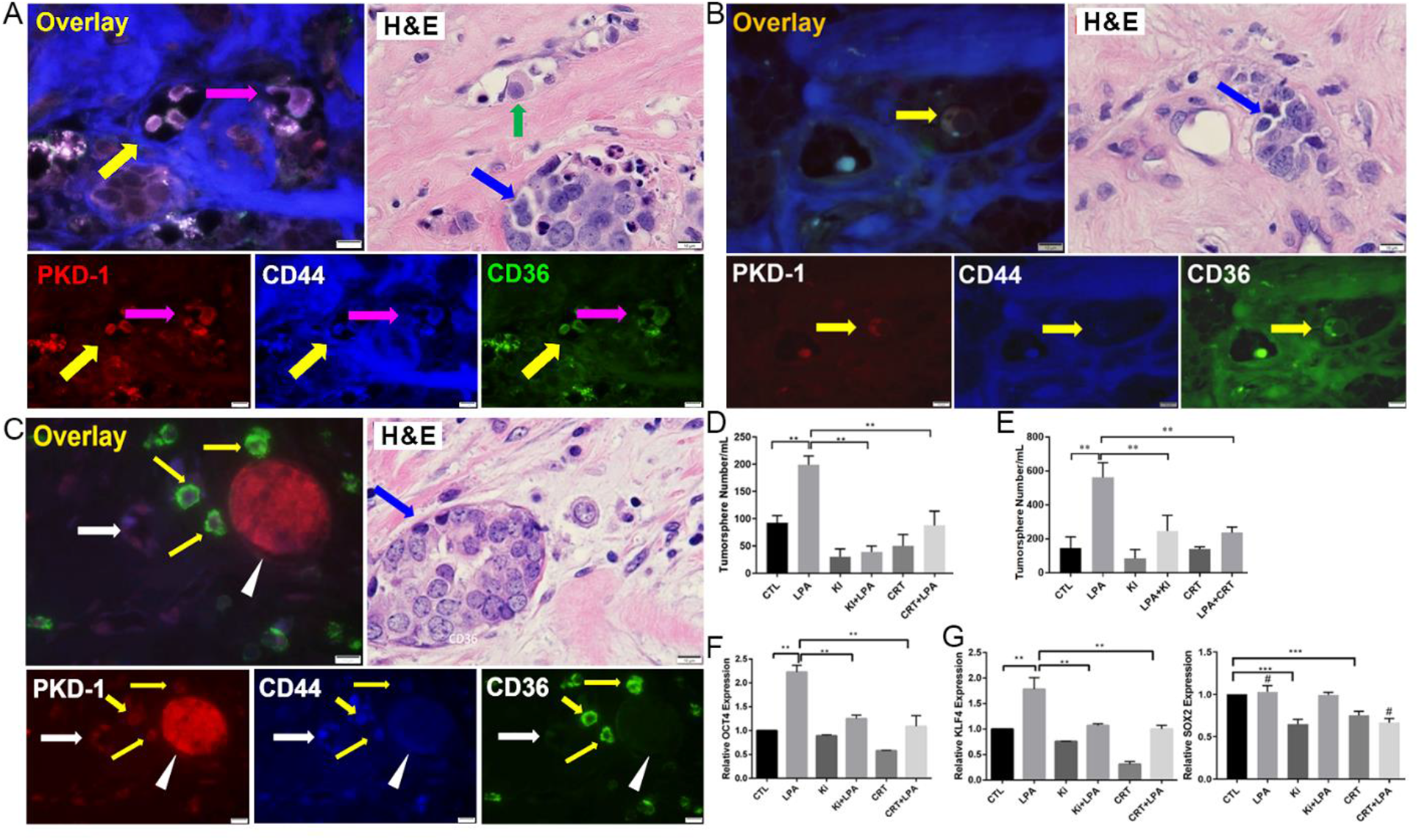
Distribution of breast cancer stem cells expressing both PKD-1 and CD36. **A.** Immunofluorescence microscopy showed that PKD-1 or CD36^+^ cancer stem-like cells (CD44^+^BCSCs) were located within the blood vessel (yellow arrows) or tumor nest (pink arrow). H & E staining as a control for tissue structure, and a tumor cell existed within the blood vessel (green arrow) and tumor nest is existed in tumor microenvironment (blue arrow). Bar = 10 μm. **B.** Immunofluorescence microscopy showed that cancer stem-like cells with positive PKD-1 or CD36 expression were located nearby the tumor blood vessels (yellow arrows). H & E staining showed tissue structure of the breast cancer and tumor nest was indicated by a blue arrow. Bar = 10 μm. **C.** A small subset of CSCs expressing moderate levels of PKD-1 but high levels of CD36 (yellow arrow) were located outside of cancer nest that expressed high levels of PKD-1 but moderate levels of CD44 (white arrow head) in human BC. A small subset of CSCs expressed moderate levels of PKD-1 and CD44 but minimal levels of CD36 and appeared to form vessel-like structure (white arrow). Blue arrow in the H & E indicates tumor nest. Bar = 10 μm. **D.** Tumorsphere formation of mouse BC cells. CD44^+^ E0771 cells were cultured in complete MammoCult™ Medium with the treatment of 10 *μM* LPA, 1 *μM* CRT0066101 (PKD inhibitor), 1 *μM* Ki16425 or their combination for 7 days. **E.** T umor sphere formation of ER^+^BC cells. Human BC (MCF-7) cells were cultured in complete MammoCult™ Medium with the treatment of 10 *μM* LPA, 1 *μM* CRT0066101 (PKD inhibitor), 1 *μM* Ki16425 or their combination for 7 days. The mammary spheres were counted under the OLYMPUS CK30 microscope, triplicate experiments were performed, and the results are shown as the mean value ± SEM. ** *P* < 0.01. **F.** LPA-PKD-1 signaling stimulated expression of Oct4 and KLF4 in ER^+^BC cells. **G.** Sox2 expression was regulated differently by LPA/PKD-1 signaling. Total RNA was extracted from MCF-7 tumorspheres with different treatments, and mRNA levels were assayed with RT-qPCR. The results of triplicate experiments are shown as mean ± SEM. \*\**P* < 0.01, \*\*\**P* < 0.001 or #*P* < 0.01.

LPA is known to function via activation of PKD-1 pathway in ECs and cancer (34, 72, 73) and LPA/PKD-1 signaling may be associated with stemness of cancer cells (37–39). We next assessed whether the LPA/PKD-1 signaling is critical for the maintenance and self-renewal of BCSCs by using mammosphere formation assays that can identify functional BCSCs *in vitro* (74).

By taking advantage of the anchorage-independent property of BCSCs for cell expansion and enrichment (2, 75) in ultra-low attachment plates, we initially optimized the growth conditions to observe the role of LPA/PKD-1 signaling in BCSCs grown as mammospheres. After optimizing growth conditions, we reproducibly showed that LPA/PKD-1 signaling significantly promoted the self-renewal of BCSCs in both mouse (Figure 4D) and human (Figure 4E) BC cells. To define the association between LPA/PKD-1 signaling and BCSCs, we analyzed changes in the expression of stemness-associated genes in mammospheres treated with either control, LPA, or PKD inhibitor (CRT). BCSCs exposed to LPA significantly increased the expression of *Oct4* and *KLF4* (Figure 4F), genes that sustain self-renewal capacity (76, 77). Addition of either a LPA antagonist or a PKD inhibitor attenuated the LPA-induced expression of *Oct4* and *KLF4* (Figure 4F).

Unexpectedly, LPA treatment did not increase the expression of *Sox2,* a transcription factor that is expressed in CSCs and mediates resistance towards established cancer therapies (78) (Figure 4F). However, inhibition of endogenous LPA/PKD-1 signaling by either LPA antagonist or PKD inhibitor decreased the gene expression of Sox2. Furthermore, LPA exposure recovered LPA antagonist-mediated, but not PKD inhibitor-mediated, Sox2 downregulation (Figure 4G).

To validate the role of LPA/PKD-1 signaling in supporting BCSC stemness, we evaluated the impact of PKD-1 depletion on BCSC tumor initiating potential in well-established in vitro limiting dilution tumor assays. Transfection of siRNA demonstrated an efficient knockdown of endogenous PKD-1 expression at protein level (Figure 5A). Furthermore, knocking down PKD-1 expression significantly impaired the frequency of repopulating of the cells when compared with the control (Figure 5B, C & D). These results suggest that LPA/PKD-1 signaling may be critical in BCSC maintenance and expansion as well as tumor initiation capacity.

**Figure 5.**
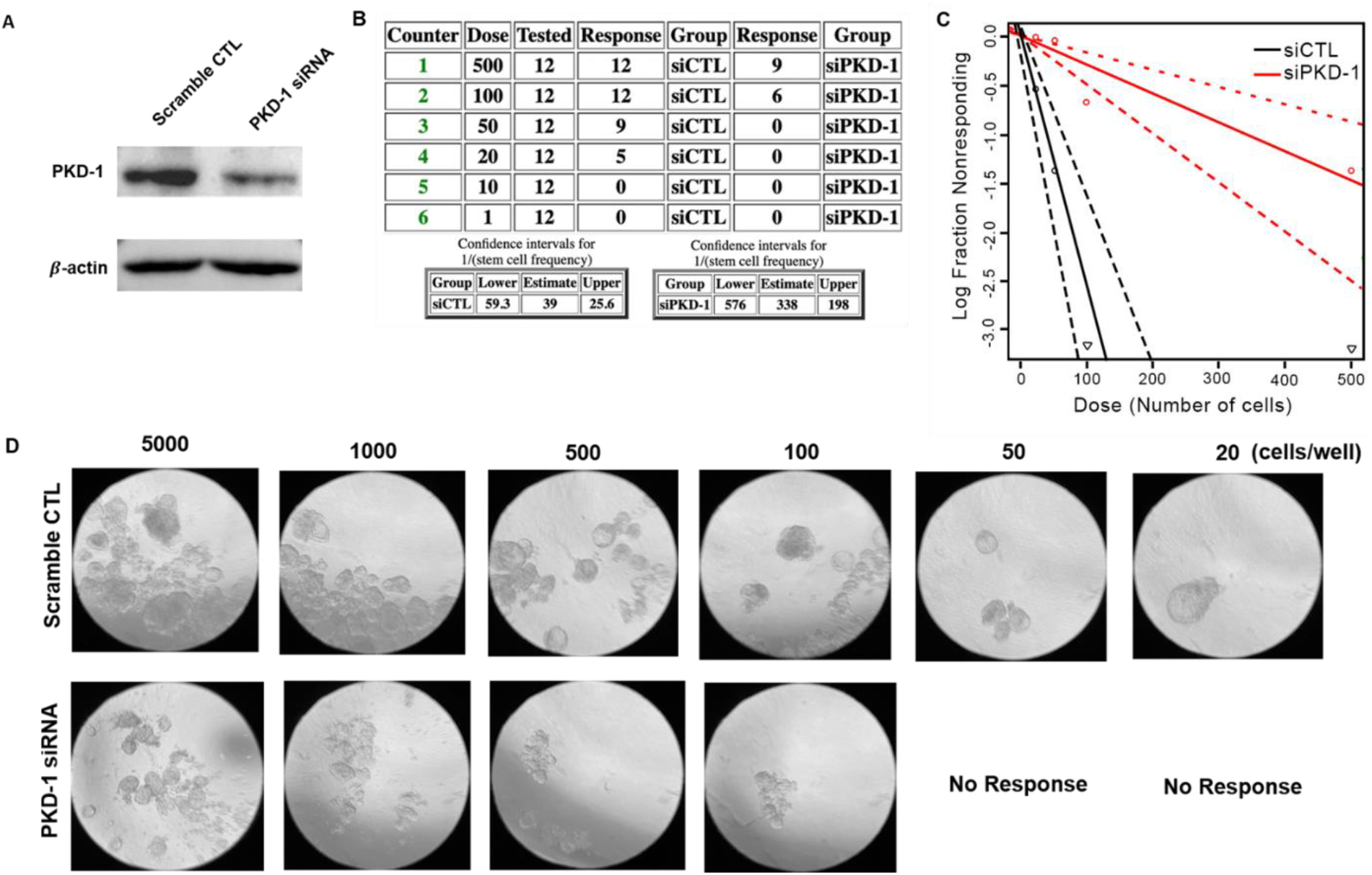
PKD-1 is required for tumorigenicity by tumorsphere formation efficiency assay. **A.** Immunoblotting assay indicates that the PKD-1 expression decreased when transfecting MCF-7 with PKD-1 siRNA as compared with the scramble control (CTL). **B.** MCF-7 with PKD-1 depletion significantly reduced tumorsphere formation efficiency. **C.** Quantitation graph shows that the number of tumorsphere formed in each well with the seeding density starting from 500 to 20 cells/well by statistical analysis. **D.** Representative images for tumorsphere formation in MCF-7 cells subjected to siRNA scramble control (CTL) and PKD-1 siRNA and with different seeding density.

LPA treatment of human BC cells and BC-bearing mice suggested that PKD-1 signaling likely increased the expression of ALDH1A1 (Figure 6A). To further define the role of the LPA/PKD-1 signaling pathway in BC progression, we exposed human BC cells to either LPA, LPA antagonist, or a PKD inhibitor to examine changes in ALDH1A1 expression. Immunofluorescence microscopy showed that the expression of ALDH1A1 was significantly increased in response to LPA/PKD-1 signaling (Figure 6B). Given that Notch1 activity plays a pivotal role in the stemness and progression of ER^+^ BC (79, 80) and ECs can interact with BC via the Notch1 pathway (10), we tested whether PKD-1 signaling can impact Notch1 expression by transducing BC cells with PKD-1. Overexpression of PKD-1 significantly increased the expression of Notch1 at both the mRNA and protein level in BC cells (Figure 6C & D), along with the increased expression of CD44 and KLF4 (Figure 6E & F). To provide a mechanistic explanation for LPA-induced BC stemness, we examined the downstream of PKD-1 signaling MAPK/Erk1/2. The results showed that LPA-mediated MAPK/Erk1/2 phosphorylation was inhibited by a PKD inhibitor (Figure 6G), suggesting that LPA/PKD-1 signaling may stimulate BC stemness via MAPK/Erk1/2-mediated stemness-associated gene transcription.

**Figure 6.**
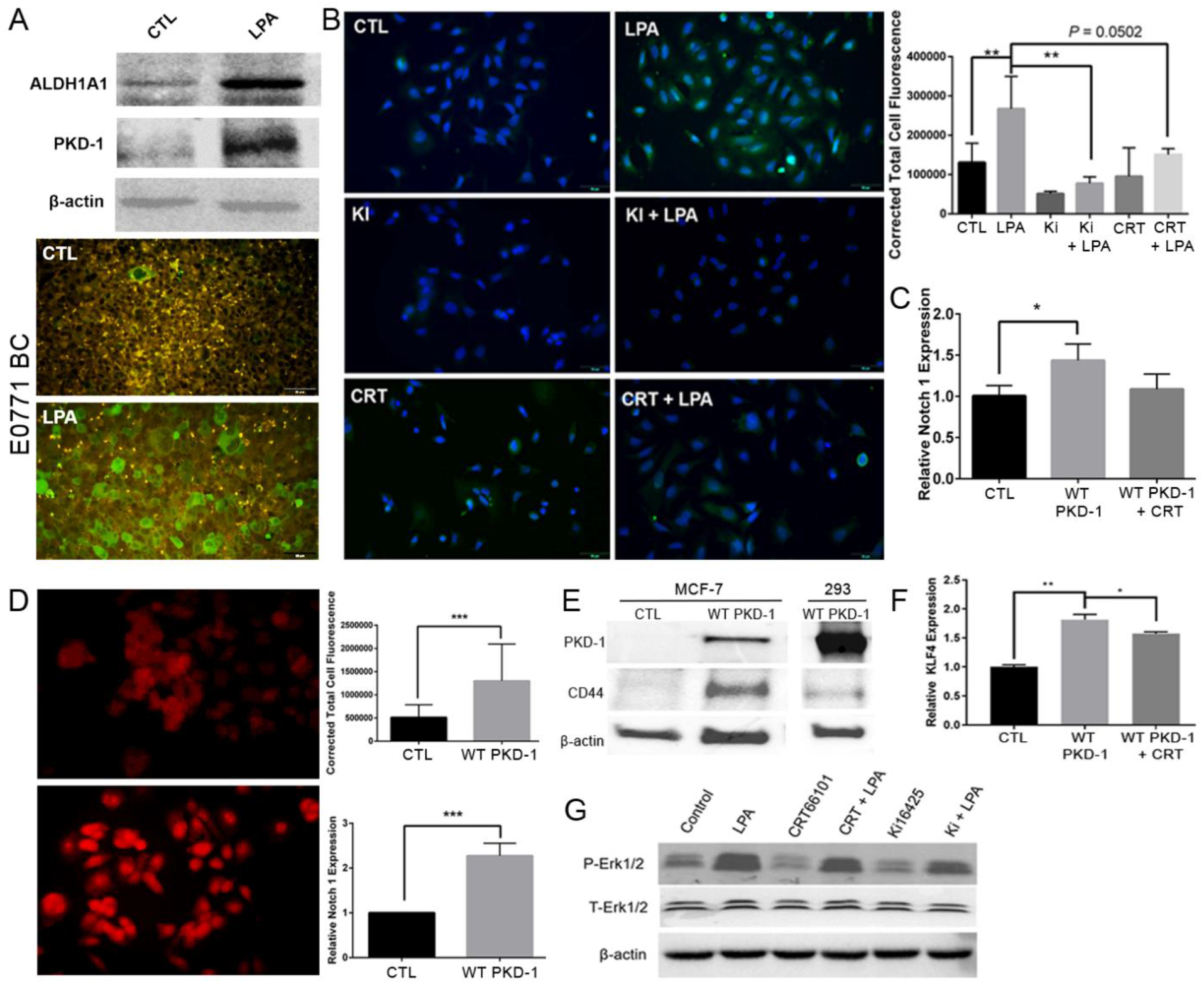
LPA-PKD-1 signaling pathway in self-renewal and plasticity of breast cancer stem cells. **A.** Induction of ALDH1A1 expression in BC cells (upper panel). The E0771 cell were cultured in RPMI-1640 medium with 10% FBS and 1% Penicillin/Streptomycin. Following starvation in serum free medium for 6 hours, the cells were treated with LPA (10μM) for 24h, the cell lysates were collected for Western Blot. Duplicated experiments were performed, and shown is a representative result; ALDH1A1 expression in implanted E0771 BC (lower panel). Immunofluorescence microscopy was used to detect ALDH1A1^+^BC cells in tumor tissues. Representative images are shown. Bar = 50 μm. **B.** MCF-7 cells were exposed to 10 *μM* LPA, 1 *μM* CRT0066101 (PKD inhibitor), 1 μM Ki16425 (LPA1,3 antagonist), and their combinations for 24 hours. The control and treated MCF-7 cells were incubated with ALDH1A1 antibody followed by appropriate secondary antibody. Representative images are shown from triplicate experiments. Fluorescence intensity was measured by ImageJ and calculated by the corrected total cell fluorescence (CTCF). The relative expression was shown as the mean ± SEM. \*\**P* < 0.01. **C.** Total RNA was extracted from the MCF-7 control and those transduced with PKD-WT and treated with PKD inhibitor CRT0066101. The mRNA levels were assayed by RT-qPCR. \**P* < 0.05. **D.** MCF-7 cells transduced wild type PKD-1 (PKD-WT) were stained with Notch 1 antibody, followed by appropriate secondary antibody. The fluorescence intensity was measured by ImageJ and calculated by the CTCF. The mRNA levels were assayed with RT-qPCR. Triplicate experiments are performed and results shown as the mean ± SEM. \*\*\**P* < 0.001. **E.** Overexpression of wild-type PKD-1 increased CD44 expression in BC cells. MCF7 cells were transduced with PKD-WT and protein lysate was collected for Western blots. MCF7 without PKD-WT or 293T cells transduced with PKD-WT as controls. **F.** Overexpression of PKD-WT induced KLF4 expression in BC cells. MCF7 cells transduced with PKD-WT were starved in serum free DMEM media overnight, and treated with/without CRT0066101 (0.5 μM) for 24 hours. Total RNA was extracted for detection of KLF4 by RT-qPCR. \**P* < 0.05 or \*\**P* < 0.01. **G.** MCF7 cells were serum-free starved overnight and then treated with LPA (10 μM) and/or LPA antagonist Ki16425 (1 μM) or CRT0066101 (1 μM) for 24h. The cell lysates were collected for Western Blot. Duplicate experiments were performed.

### Discussion

CSCs are intimately connected with the vasculature and contribute to tumor growth, recurrence and metastasis. They can accumulate in the perivascular regions in tumor microenvironment (32, 81). The interactions between CSCs and their heterogeneous vasculature (21,22) may determine CSC development and fate. Additionally, circulating tumor cells contain a distinct subset of circulating CSCs that bear metastasis-initiating capabilities (44), and more CSCs exist in arterial blood than in venous blood (82). The arteriole-rich endosteal regions in bone marrow microvascular compartment show the highest partial oxygen pressure and contain hematopoietic SCs (83, 84). The hematopoietic SCs appear near nestin-positive arteries in bone marrows (84). Furthermore, arterialization may contribute to liver and lung metastases (28, 29). These studies strongly suggest that arteriolar niche contributes to the maintenance and expansion of CSCs. Consistent with these observations, our study highlights the existence of the arteriolar niche in the tumor microenvironment of ER^+^BCs. Moreover, BCSCs tend to be enriched in the arteriolar niche. Arteriolar ECs may crosstalk with BCSCs via Notch signaling pathway, which likely links to LPA/PKD-1 signaling-mediated arteriolar differentiation. Importantly, our finding that aggressive CSCs tend to be enriched in the arteriolar niche, which may enjoy a high level of perfusion, are inconsistent with the notion that tumor hypoxia is a major contributor of tumor aggressiveness. Therefore, our study uncovers a previously underappreciated vascular type within the tumor microenvironment to promote BC progression likely via perfusion of nutrients and oxygen. This finding is supported by a recent seminal study (85) and other previous studies (22).

The crosstalk between tumor cells and their microenvironment is crucial for cancer cell selfrenewal, tumor growth and metastasis (12, 32). Notch1 can mediate cell–cell communication in many tissues (48, 86). Endothelial Notch signaling may promote tumor progression and SC phenotype including self-renewal of CSCs (10, 87–90). In triple negative BC, ECs can enhance self-renewal, survival, and pro-metastatic properties of BCSCs through direct cell-cell contact via the Notch signaling in vitro (10). Similar to these studies, we demonstrate that co-culture of ER^+^ BC cells with HMVECs with high levels of DLL4 increases stemness-associated gene transcription in cancer cells including *CD44, ALDH1A1, KLF4,* and *CD36,* along with increased *PKD-1* expression, suggesting the stimulating effect of arteriolar Notch signals to regulate the stemness via the PKD-1 pathway. LPA/PKD-1 signaling could mediate crosstalk between arteriolar ECs and BCSCs, and this may provide insights into the role of arteriolar niche in CSC plasticity, and pro-tumoral and pro-metastatic properties as ECs can directly interact with BCs, and indirctly stimulate CD44 and ALDH1A1 expression (40, 42, 91, 92).

Furthermore, in BC cells PKD-1 appears to mediate long-term signaling responses as shown by LPA-stimulated sustained-phosphorylation of MAPK/Erk1/2. This may be PKC-independent (93) and involved in the regulation of stemness-associated gene transcription. However, much remains to be investigated as to how the extracellular signals regulate these interactions and what additional diffusible signals is released from the arteriolar vasculature to promote stemness directly and indirectly within the tumor microenvironment.

Previous studies suggest that the LPA/PKD-1 signaling stimulates arteriolar differentiation, along with a decrease in CD36 and increase in DLL4 expression in MVECs (30, 31, 94), and promotes ER^+^ BC progression (30, 31, 33, 69). We now show that PKD-1 deficiency in lung ECs upregulates the expression CD36 but downregulates arteriogenic gene expression. This suggests that the LPA/PKD-1 signaling axis is required for proarteriogenic reprogramming for arteriolar differentiation, thereby likely enhancing the crosstalk between ECs and CSCs (10, 95). Although distinct vascular niches are likely needed to regulate BCSCs, the LPA/PKD-1 signaling-mediated DLL4 expression in the arteriolar ECs facilitates direct EC interaction with Notch1^+^ stem-like cells. This may be an additional mechanism by which vascular niche promotes maintenance and expansion of CSCs within the tumor microenvironment.

In fact, EC sprouting may abolish inhibitory growth signals in dormant tumor cells to allow metastasis of BC cells in different organs (9). Additionally, Notch signals from the stromal niche can induce the reactivation of growth and self-renewal pathways (96). Therefore, direct endothelial DLL4-mediated Notch activation in circulating BCSCs in arteriolar niche could promote survival and metastatic potential of the cancer cells. TAECs with arteriolar phenotype may directly or indirectly provide signals that actively promote BC stemness. We thus propose a working model that PKD-1 signaling may promote BC stemness by activating stemness-related Notch pathway in cancer cells and by creating an arteriolar niche via stimulating arteriolar differentiation within the tumor microenvironment, thereby promoting the self-renewal of BSCCs. This pathway can also directly stimulate expression of Notch1, CD44 and KLF4 for the expansion of BCSCs. Furthermore, LPA/PKD-1 signaling-mediated ALDH1 expression may regulate the plasticity of BCSC whereas the increased expression of both ALDH1 and CD36 is likely significantly increase the metastatic potential via activation of LPA/PKD-1 signaling (Figure 7). Moreover, the arteriolar niche could equip and educate peri-arteriolar and circulating BCSCs to escape from immune surveillance, survive in the circulating blood, which facilitates their spread to the liver and lung. Further investigations are needed to provide a deep understanding of whether and how individual factor(s) or a combination of factors are required to maintain this unique niche.

**Figure 7.**
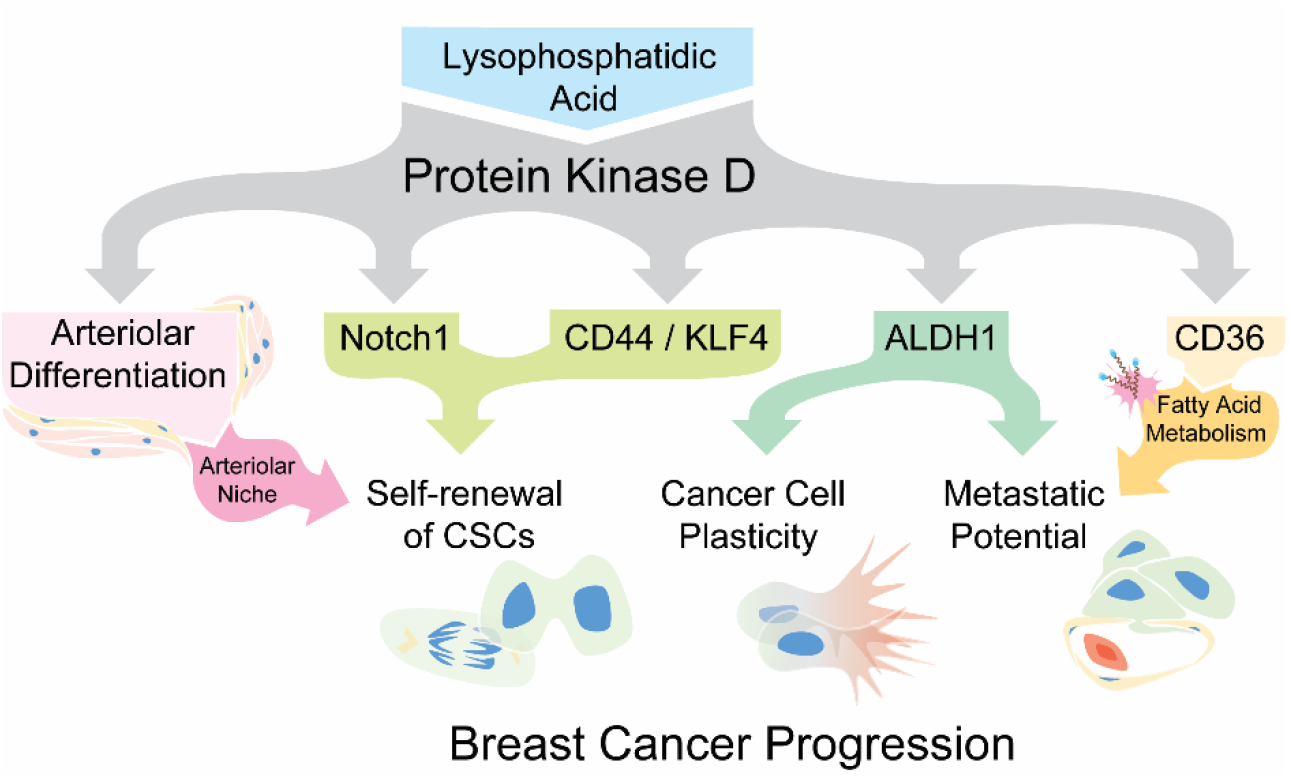
A proposed mechanistic model shows that the LPA/PKD-1 signaling regulates arteriolar differentiation within TME, CSC plasticity and BC progression.

LPA/PKD-1 signaling also contributes to BCSC specification by increasing the expression of genes encoding pluripotency factors and self-renewal of BCSCs. Using human and mouse ER^+^ BC cell lines that contains CSCs, which generate clonal variability (97) and lead to metastases (98), a syngeneic ER^+^ BC mouse model, and tumor samples from patients with ER^+^ BC, we demonstrate the positive correlation of this pathway with stemness-related gene signature. We show an increased expression in CD44, ALDH1 and CD36 and Notch1 in response to LPA/PKD-1 signaling. In current study we focus on ER^+^ BC because LPA is critical to ER^+^ BC progression (31) and PKD-1 is expressed in ER^+^ BC but not in triple negative BC (TNBC) (99, 100) and CD36 is also highly expressed in ER^+^ BC (46, 101). However, we do not exclude the possibility that a subset of TNBC cells may express PKD-1 as PKD-1 expression increases in BCSCs. It is also known that the ER^+^ BC is characterized by late metastatic relapses (102), in which CSCs may play an important role (103, 104).

Finally, CSCs may be much more plastic and abundant and can proliferate vigorously (105, 106). The plasticity of CSCs allows them to change between CSC and non-CSCs, representing a substantial divergence when compared to normal SCs (107). CD44 and ALDH1A1 are molecular markers for BCSCs and present BC stem-like features including high tumor-initiating and self-renewing capabilities as well as metastasis and progression (2, 40–44). Furthermore, BC cells with expression of both CD44 and ALDH1A1 show the highest degrees of plasticity (40, 42, 91, 92), in which the LPA/PKD-1 signaling may play an essential role. We speculate that this pathway-mediated plasticity may be determined by cues from proangiogenic and proarteriogenic TAECs, thereby conferring tumors with aggressive and lethal properties including drug-resistance and metastatic dissemination. Targeting this pathway may have additional therapeutic benefits by counteracting acquired resistance via eradication of ER^+^ BCSCs (108). Not only are the BCSCs enriched in arteriolar niche, intriguingly, this study also shows that the BCSCs that are positive for both PKD-1 and CD36 come out of tumor nests and appear within the vascular niche, particularly arteriolar niche. Additionally, PKD-1 signaling may promote metastatic potential of BCSCs via upregulation of CD36 in BCs (46), and LPA signaling was also proposed as a therapeutic target for vascular inflammation, atherosclerosis, and potentially calcific aortic valve disease (109). Our current study indicates that inhibition of the LPA/PKD-1 pathway may likely disrupt the arteriolar niche for CSCs and eradicate BCSCs as a double-edge sword, thereby alleviating therapeutic resistance and controlling relapse and metastasis of cancers. Therefore, LPA/PKD-1 signaling regulation of arteriolar differentiation and stemness features of BCSCs highlights the clinical relevance and should provide insights into therapeutic strategies against BCs and such other types of highly angiogenic cancers as lung cancer, melanoma and pancreatic neuroendocrine cancers.

## MATERIALS AND METHODS

### Study approval

All *in vivo* experiments were conducted in accordance with the Guide for the Care and Use of Laboratory Animals of the NIH. Animal studies were conducted under approved IACUC from University of Alabama at Birmingham and Medical College of Wisconsin. Tumor specimens from human patients with ER^+^ breast cancer were used to perform immunohistochemistry and immunofluorescence experiments without any link to subject identifiable information.

### Reagents and antibodies

Oleoyl-L-α-lysophosphatidic acid (LPA)(L7260) was purchased from Sigma-Aldrich. The PKD inhibitor CRT0066101 (A8679) was purchased from APExBio. The LPA inhibitor Ki16425 (10012659) was purchased from Cayman Chemical Company. Gelatin (9000-70-8) was purchased from Bio-Rad. The reagents for RT-qPCR include the RNeasy Mini Kit (Qiagen), Power SYBR Green PCR Master Mix and High-Capacity cDNA Reverse Transcription Kit (Applied Biosystems). RT^2^ qPCR Primer Assay and primers for human/mouse CD36, EphrinB2, Neurophilin 1, DLL4, PRKD-1, Oct4, Sox2, and GAPDH (Qiagne and IDT). Antibodies include PKD/PKCμ (D4J1N), Neurophilin1 (D62C7), phospho-Erk (1/2) (D13.14.4E), and human/mouse Erk (1/2) (137F5) (Cell Signaling Technology), ALDH1A1 (AF5869) (R&D Systems) and EphrinB2 (ab131536) and CD36 (ab133625) (Abcam). Antibodies for immunostaining include rabbit anti-human/mouse PKD-1 (4502371) and mouse anti-mouse α-smooth muscle actin(A2547) (Sigma-Aldrich), mouse anti-human/rat CD44 (5640S) and rabbit anti-human/mouse ALDH1A1 (12035S) (Cell Signaling Technology), rabbit anti-human/mouse CD36 (ab133625) (Abcam), AlexaFluor 594 conjugated donkey anti-rabbit IgG (A21207), AlexaFluor 594 conjugated donkey anti-mouse IgG(A21203), AlexaFluor 594 conjugated donkey anti-goat IgG(A11058), AlexaFluor 488 conjugated donkey anti-goat IgG(A11055), AlexaFluor 488 conjugated donkey anti-rabbit IgG(A21206), and AlexaFluor 488 conjugated goat anti-mouse IgG(A11001) (Invitrogen). VECTASHIELD Antifade Mounting Medium with DAPI (H-1200) was purchased from VECTASHIELD. Transfection reagent TransIT-X2 reagent (MIR6003) was purchased from Mirus. Opti-MEM I reduced-serum medium (51985–034) was from Gibco.

### Cell culture

Human microvascular endothelial cell line (HMVECi-D)(94) were grown in endothelial media (DMEM medium with 5% MVGS and 5% FBS) under 5% CO_2_, 37°C. Primary mouse lung endothelial cells were isolated as previously reported (34, 58) and cultured as in HMVECi-D. E0771 cells were cultured as described previously (31). MCF7 cells were seeded in T75 flasks or 10 cm plates in low glucose Dulbecco’s modified Eagle’s medium (DMEM) containing 10% fetal bovine serum (FBS), 2 mM glutamine, 0.01 mg/ml insulin and 1% penicillin/streptomycin mix and incubated at 37°C in an atmosphere of 5% CO_2_. A co-culture of MCF-7 cells transduced with Ds-Red and HMVECi-Ds were plated onto glass bottom dishes at the ratio of 2:1 for 48 hours for cell-to-cell contact analysis.

### Isolation and culture of primary mouse lung endothelial cells

Isolation and culture of lung endothelial cells from control and endothelial-specific *pkd-1* knock out mice were modified in reference to a previous study (110). Briefly, lungs were collected from 3 to 4 adult mice and minced, digested with pre-warmed collagenase (Sigma C0130) at 1 mg/ml in DPBS. After incubation at 37°C for 45 min, tissue suspension was pipetted through a 70 μm cell strainer. The suspension was then spin down at 400 g for 4 min at 4°C. The pellet was resuspended pellet in 1ml of cold PBS with 0.1%BSA by adding 30ul of Dynabeads (Invitrogen, Cat#11035) conjugated with rat anti-mouse CD31 antibody (BD, Cat# 553370) and incubated on a rotator at room temperature for 10min. Subsequently, the beads with cells were separated in a magnetic separator, along with five washes in cold PBS+ 0.1%BSA. Finally, assumed CD31 ^+^ endothelial cells were re-suspended in DMEM media with 5% MVGS (Gibco, Cat# S00525) and 5% FBS and seeded in a gelatin-coated dish.

### Transwell endothelial cell and breast cancer co-culture

Co-culture of endothelial cells with cancer cells without cell-to-cell contact was referred to a previous study ^(87)^ and performed in the Transwell cell culture chamber (Corning) inserts with 0.4 μm pores and six-well plates. MCF-7 and HMVECi-D were cultured in DMEM (Corning) with 10% Fetal Bovine Serum and 1% Penicillin/Streptomycin and EGM-2 Medium (EBM-2 Basal Medium with EGM^TM^-2 MV Microvascular Endothelial Cell Growth Growth Factor, Lonza) respectively for 18 hours before starting the co-culture of the two cell lines. MCF-7 was seeded in the 6-well plate while the HMVECi-D was cultured in the insert with transparent PET membrane (0.4 μm pore size, Corning) with a ratio of 5:1. 6×10^4^ cells were seeded in the insert while 3×10^5^ cells grown in the well. HMVECi-D medium was switched to the MCF-7 culture medium when the two cells were co-cultured for 72 hours. They were co-cultured in MCF-7 media for 48 hours. Total RNA of MCF-7 cells was then extracted and assayed by RT-qPCR.

### Plasmid transduction and transfection

Lentiviral system was used to transduce genes into E0771 BC cells, mouse BC stem-like cell (CSC) line derived from E0771 and HMVECs. Five to ten MOI of lentiviral particles containing either luciferase: GFP or Ds-Red or wild type PKD-1 (PKD-WT) were added to the E0771 or HMVECi-D for the transduction. An EVOS^®^FL cell imaging system or flow cytometry was used to determine transduction efficiency. For plasmid transfection, MCF-7 cells with high PKD-1 expression were incubated in a 6-well plate with a seeding density of 6.0×10^5^ cells/well for 24 hours. The scramble control and PKD-1 siRNA (IDT) was then respectively transfected into the cells in Opti-MEM I reduced-serum medium (Gibco) by TransIT-X2 transfection reagent (Mirus) for 6-18 hours and replaced with BC culture media containing 10% FBS for an additional 48 hours.

### Real time quantitative RT-qPCR

Gene expression was assessed by real-time quantitative RT-qPCR as previously described (*1*). The RT2 qPCR primer assays (Qiagen) or qPCR primers (IDT) for the target genes and housekeeping genes were used for PCR reactions. Total RNA was isolated from breast cancer cells or ECs using the RNeasy Mini Kit (Qiagen) and then subjected to real-time qRT-PCR using CFX Connect Real-Time System (Bio-Rad). The following genes were assayed with GAPDH or PPIA transcripts that were amplified in separate wells for normalization of variances in input RNA. The relative Ct value was used to compare the fold or quantitative change of mRNA expression.

### Immunoblot assays

Immunoblots of cell lysates were probed with relevant antibodies. Protein concentrations were assayed with a BCA kit (Pierce Chemical) and β-actin was used as loading control. Cell lysates were separated with commercially ready (Bio-Rad & Fisher) or self-made gel and subjected to Western blots. Densitometry was performed using NIH Image J.

### Immunofluorescence and immunohistochemical assays

Tissues were fixed in 10% formalin for paraffin block preparation, sectioned, and processed for immunohistochemical and immunofluorescence staining as previously described (34, 58). Immunofluorescence microscopy for cultured cells was performed in reference to our previous studies (30, 31).

### Mammosphere formation assays

Mammospheres assays were carried out in six well plates. A mixture of MammoCult™ Proliferation Supplement to MammoCult™ Basal Medium were prepared according to the manufacturer’s instruction. Single cells were plated in ultralow attachment plates (Corning) at a density of 2,500 viable cells in 2 ml per well. After 7 days, the colonies were counted under phasecontrast microscope by two persons. Error bars represent the standard error of the mean of three replicates.

### In Vitro extreme limiting dilution and tumorsphere formation assays

MCF-7 cells transfected with scramble control or PKD-1 siRNA were seeded with a decreasing number of cells per well (5000, 1000, 500, 100, 50, 20, 10, and 1 cell) in ultra-low attached 96-well plate. The tumorspheres were cultured in MammoCult™ Medium. The number of wells with/without growth of tumorspheres was quantified after 7 days. The data was analyzed and the log-fraction figure made using software available at bioinf.wehi.edu.au/software/elda/.

### Breast Cancer Animal model

Six-week-old female C57BL/6 mice (Jackson Laboratory) were maintained on a chow diet (D12450B, 10 kcal% fat, Research Diets, Inc). Syngeneic breast cancer were grown in the subcutaneous space of mice nearby the fourth mammary pad by implanting E0771 cells (1×10^6^ cells/mouse) (111). Mice bearing tumors for about three weeks were anesthetized and sacrificed, and tumor tissues were processed for immunohistochemistry and immunofluorescence staining. For LPA treatment experiments, vehicle or LPA (1 mg/kg) was administrated abdominally three days after tumor implantation, with an injection every three days for 18 days.

### Statistics

Quantitative data are presented as mean ± SD. Data were analyzed using 2-sided unpaired t tests using a GraphPad software package. A *P* < 0.05 or < 0.01 and 0.001 was considered statistically significant or very significant.

## Acknowledgements

We appreciate technical assistance from members of Silverstein laboratory, Brad Best, Cynthia Opansky, Liuyi Dong, and Nicholas Barnekow at Blood Research Institute as well as Gloria Yuan at Ren Laboratory at UAB School of Medicine. RH is supported by UAB School of Medicine Physician Scientist Development Fellowship. BR’s work has been supported by the American Cancer Society (86-004-26; Institution Fund to BR, Medical College of Wisconsin), the American Heart Association (13SDG14800019; BR), the Ann’s Hope Foundation (FP00011709; BR), Bioengineering Surgery Collaborative Award, University of Alabama at Birmingham (BR & AP), and the National Institute of Health (HL136423; BR).

## Supplementary Figures

**SF1.**
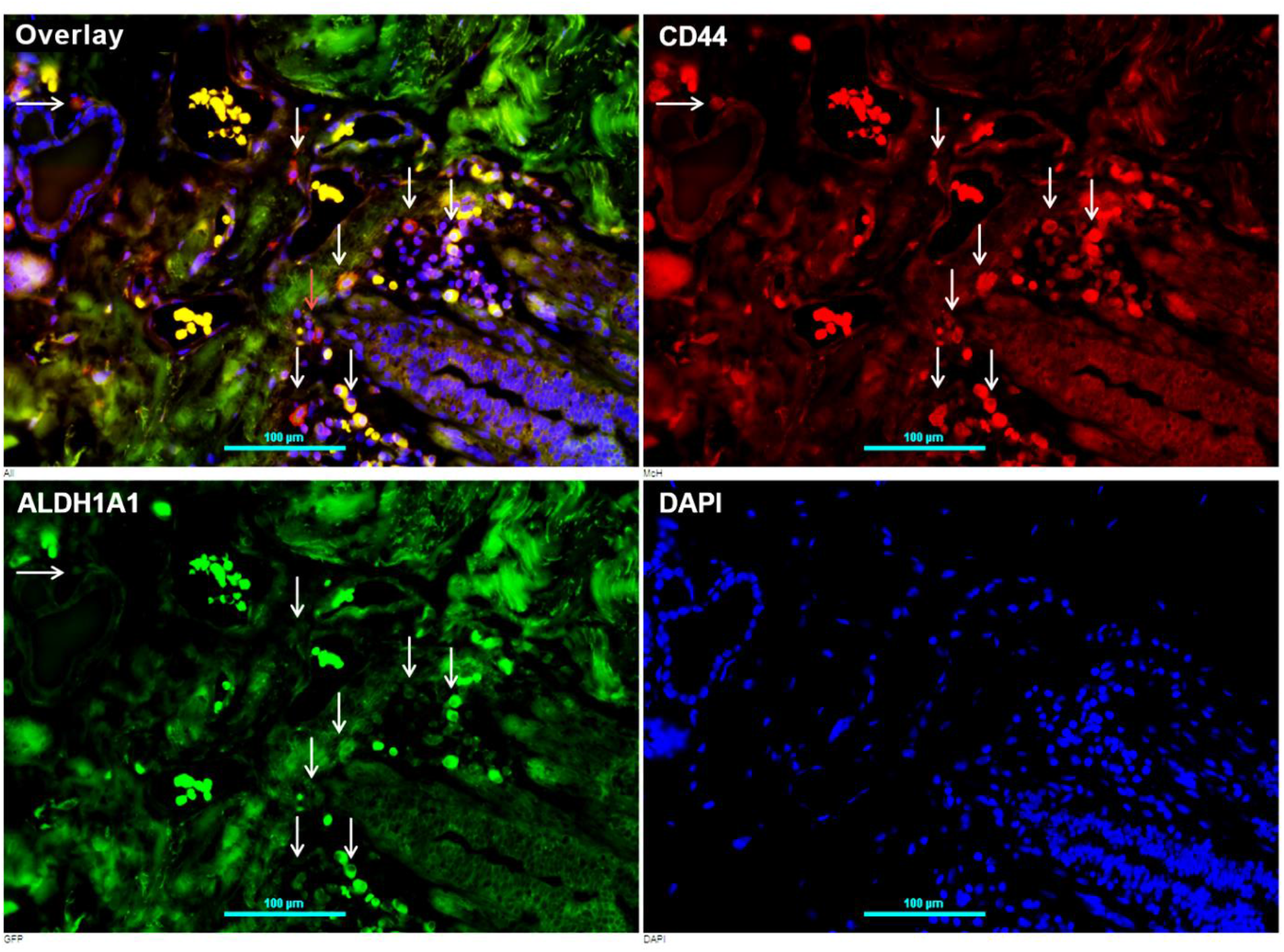
Existence of cancer stem-like cells (CSCs) in tissues from patients with ER^+^ BC. CD44 (red) and ALDH1A1 (green) positive staining in BC cells within the vascular niche (arrows); coexpression (Yellow); DAPI (blue) for the nuclei; non-specific staining in blood cells (no nuclei).

**SF2.**
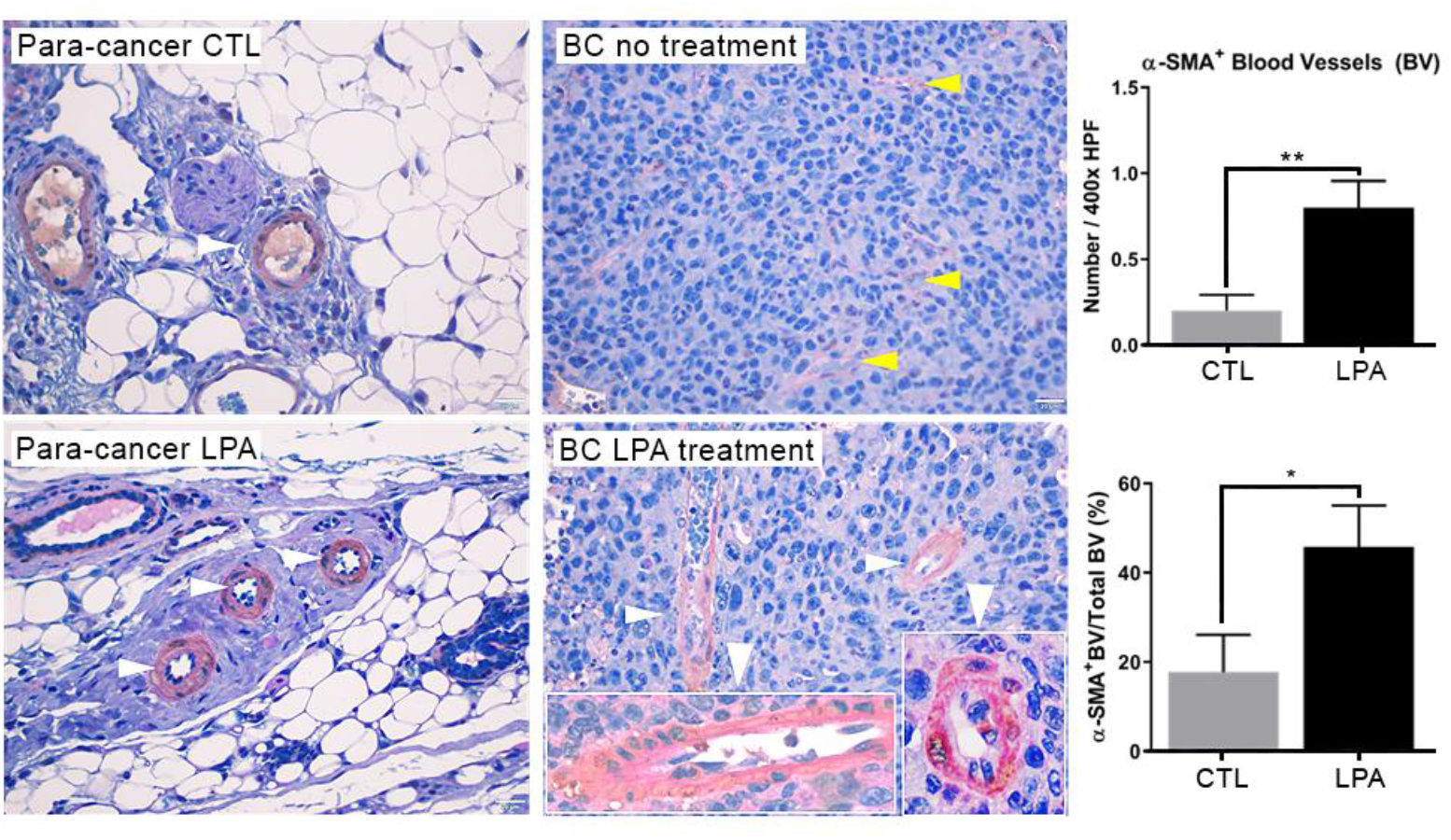
LPA treatment increased arteriolar density in BCs. Mice bearing E0771 xenografts treated with LPA (1 mg/kg) showed increased arteriolar density compared to non-treated controls as determined by number of α-SMA blood vessels (BV) and the percent of α-SMA^+^ BV per total BV. Type of BV was differentiated based on the staining and morphology. Brown and pink staining indicates α-SMA- and basement membrane-positive. White or yellow arrow heads show arterioles or capillaries, respectively. \**p*<0.05 or \*\**p* <0.01 vs control.

**SF3.**
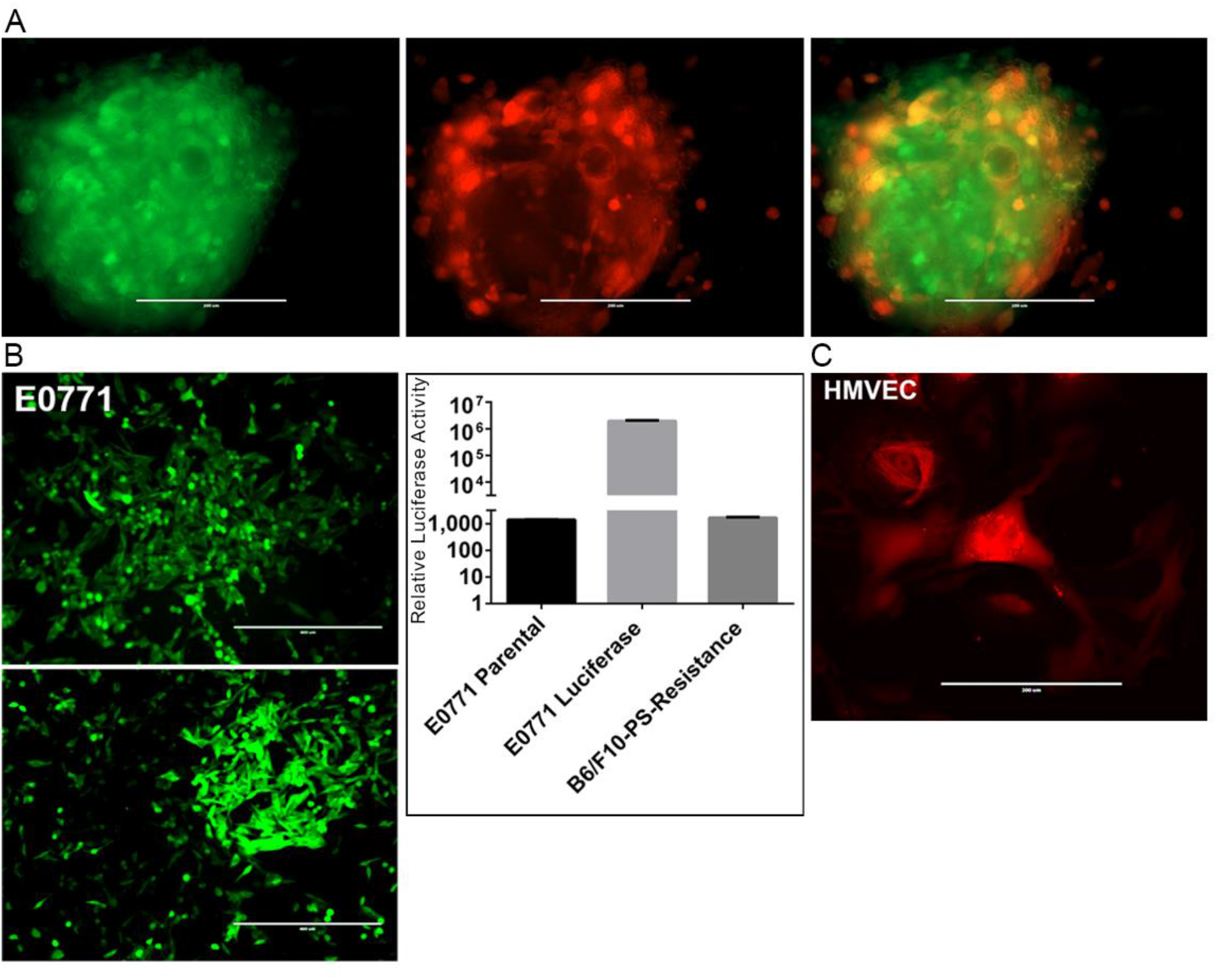
Three dimensional co-culture of BC cells and microvascular endothelial cells. **A.** E0771 cells and HMVECs were transduced with GFP or Ds-Red, respectively. Representative images are shown, scale bar = 200 μm. **B.** Representative images of E0771 cells transduced with luciferase and GFP and corresponding luciferase activities showing higher luciferase activity in E0771-transduced cells as compared to parental E0771 cells and melanoma cells (negative control), scale bar = 400 μm. GFP positive cells were sorted by flow cytometry and luciferase assays were performed using Dual-Luciferase^®^ Reporter Assay System (Promega) with a Wallac 1420 Multilabel Counter from PerkinElmer Life and Analytical Sciences (Turku, Finland). Luciferase activity is shown in the right panel. **C.** An example of HMVECs that were transduced with Ds-Red is shown.

**SF4.**
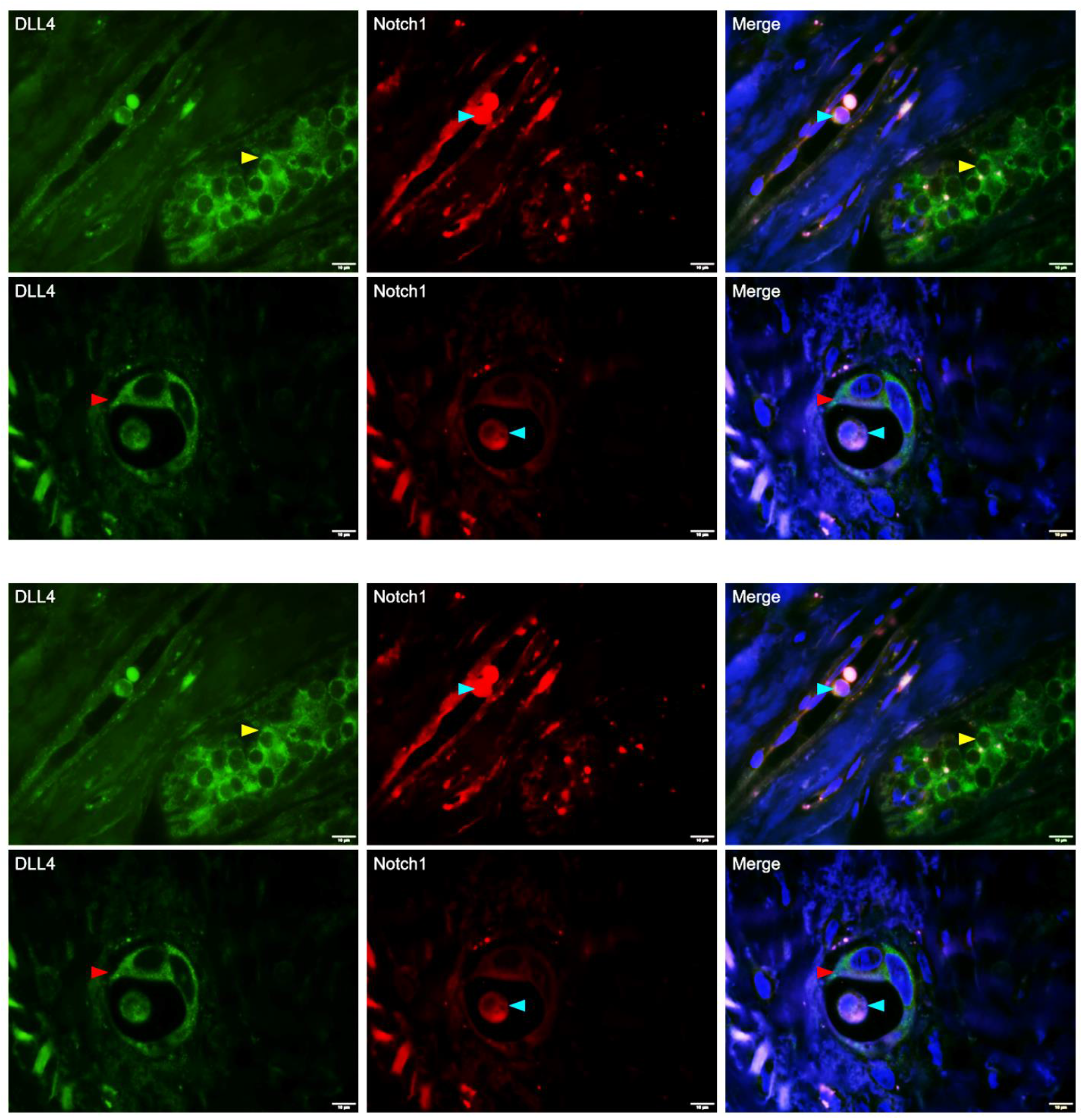
DLL-4^+^ and Notch1^+^ BC cells localized to blood vessels in patient tissues. High levels of DLL4 (stained green) were expressed in BC cells located within the cancer nest (yellow arrowheads, upper panel), while BC cells that express higher levels of Notch1 (stained red) and DLL-4 (blue arrowheads) occurred within the capillary (upper panel) or blood vessel composed of DLL-4^+^ cells (red arrowheads, lower panel). This vascular structure could be vasculogenic mimicry composed of DLL-4^+^ ECs (red arrows), which are likely derived from DLL-4^+^ BCSCs as determined by the larger size of the nucleus. Scale bar = 10 μm.

**SF5.**
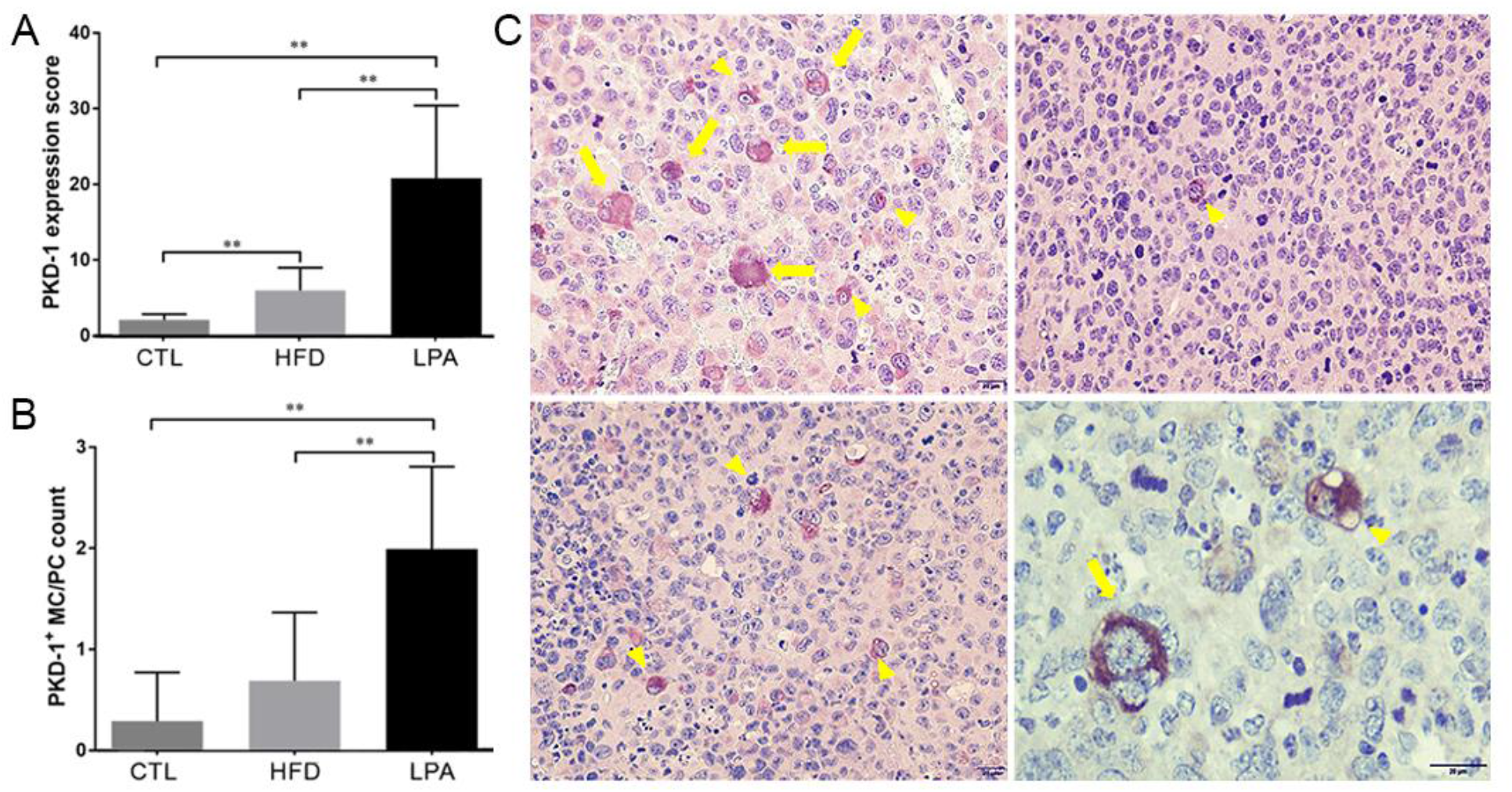
LPA treatment increased PKD-1 expression in some BC cells in an ER^+^ BC mouse model. BL6 mice were subcutaneously implanted with E0771 cells close to the mammary pads and LPA (1 mg/kg) was administered 3 days after implantation. **A.** PKD-1 expression score in a subset of BC cells in the control, LPA treatment, and high fat diet (HFD) groups \*\**p*<0.01; **B.** Higher percentage of PKD-1^+^ megakaryocyte (MC) and/or polykaryocyte (PC) per high power field (HPF) in response to LPA treatment. Shown are an average score and average number of MC and PC numbers per HPF. High positive (brown-black) cells × 2 + low positive (brown-yellow) cells×1. \*\**p*<0.01; **C.** Representative images in LPA-treated E0771 xenografts (top left), diet-induced obesity (top right), and lean control mice (bottom left), and representative magnified image of megakaryocytes and polykaryocyte (bottom right). Arrows and arrow heads denote polykaryocyte and megakaryocyte, respectively. Scale bar = 20 μm.

**SF6.**
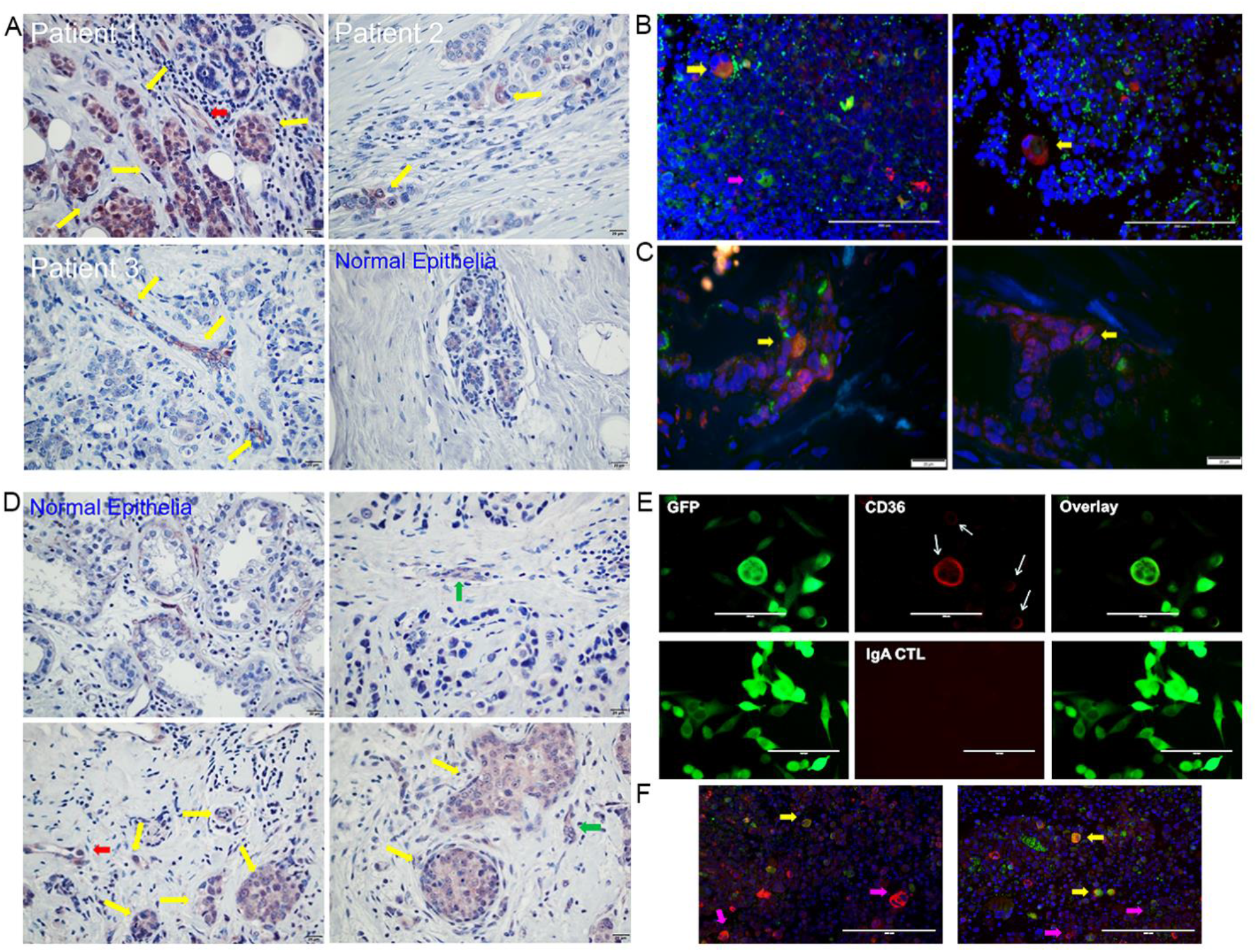
A subpopulation of breast cancer stem-like cells positive for PKD-1 and CD36 in tissues from human ER^+^ BC patients and a mouse BC model. **A.** Representative images of the population distribution of clustered or individual PKD-1^+^ cells in ER^+^ BC tissues, including clusters of BC cells with high levels of PKD-1 expression (upper left panel, yellow arrows), and individual BC cells with moderate PKD-1 expression (upper right and lower left panels). A blood vessel consisting of cells with moderate PKD-1 expression (upper left panel, red arrow). PKD-1 is weakly expressed in normal epithelia (lower right panel). **B.** Representative images of E0771 syngeneic BC tissues positively expressing CD44 (green) and PKD-1 (red); nuclei stained with DAPI (blue). Yellow arrows indicate CD44^+^/PKD-1^+^ BC cells, pink arrow shows CD44^+^ BC cells. PKD1^+^ megakaryocytes or polykaryocytes showed positive CD44 expression. Scale bar = 200 μm. **C.** Tumor tissues from patients with ER^+^ BC showed positive expression of CD44 (green) and PKD-1 (red); nuclei stained with DAPI (blue). Bar = 20 μm. **D.** Distribution of a subpopulation of CD36^+^ cells in human ER^+^ BC, including clusters of BC cells with moderate levels of CD36 expression (upper left and right panels, yellow arrows) and individual BC cells with moderate CD36 expression (upper right and lower left panels, green arrows). A blood vessel consisted of cells with moderate CD36 expression (upper left panel, red arrow). CD36 is weakly expressed in normal epithelia (lower right panel). CD36^+^ BC cells invaded into the stroma and appeared within vascular niche, presenting mesenchymal phenotype with spindle morphology (green arrows). **E.** E0771 BC cells transduced with GFP stained for CD36 expression (red). White arrows indicate the CD36^+^ cells. Representative images are shown, scale bar = 100 μm. **F.** PKD-1 and CD36 are differentially expressed in different E0771 xenograft BC cells with a small subpopulation of cells positively co-expressing both PKD-1 and CD36 (yellow arrows). The cells that expressed either PKD-1 (red) or CD36 (green) were indicated by pink arrows, nuclei stained with DAPI (blue). Representative images are shown, scale bar = 200 μm.

**SF7.**
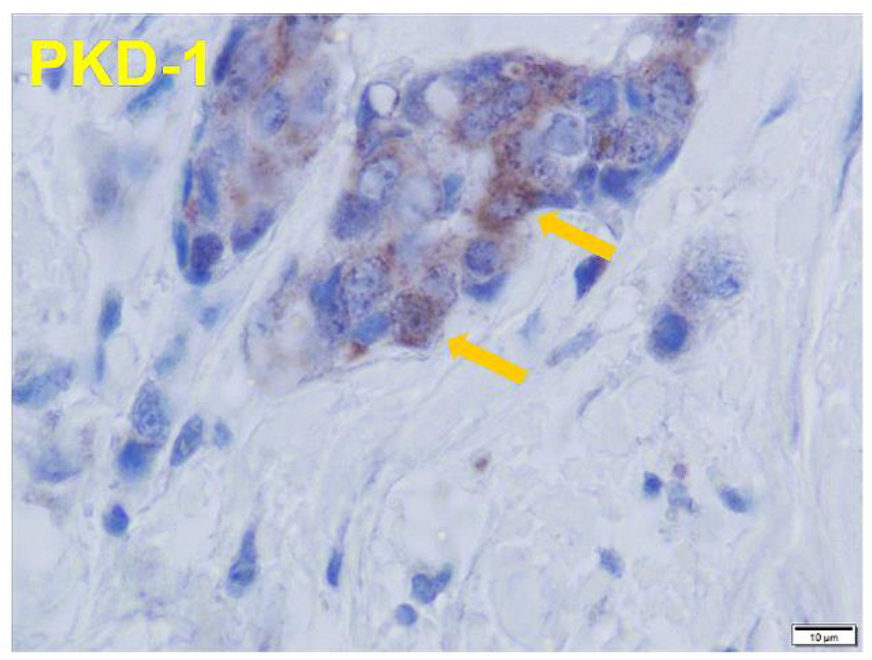
PKD-1 is highly expressed in a subset of BC cells within the cancer nest (yellow arrows).

**SF8.**
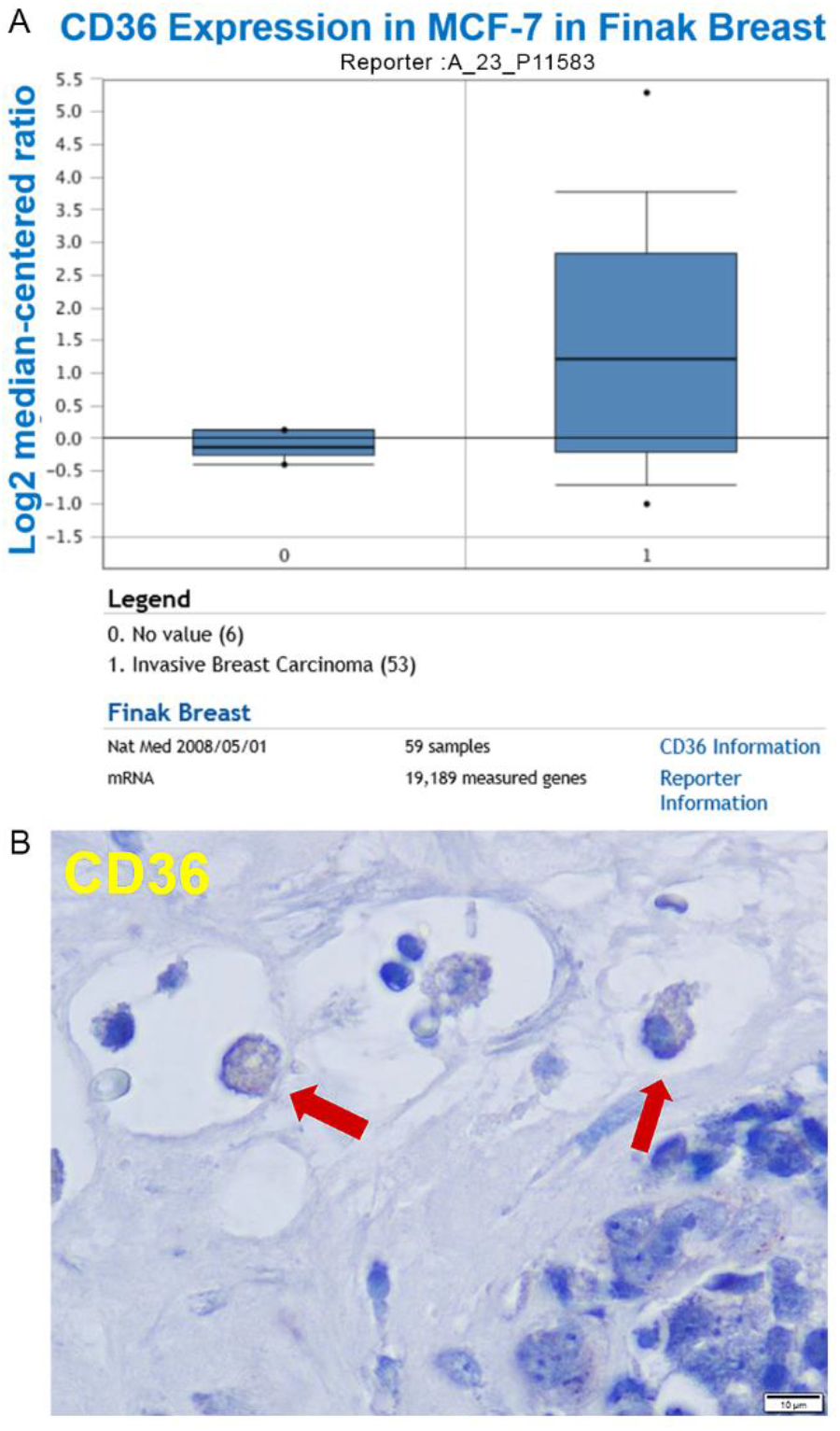
**A.** Oncomine data analysis showed that CD36 expression increased in invasive ER^+^ BC cancer compared with the control. The data were retrieved from the web link for Oncomine, a Cancer Microarray Database and Integrated Data-Mining Platform: https://www.oncomine.org/resource/login.html. **B.** Representative image of CD36^+^ expression in ER^+^ BC tissue. BC with moderate CD36 expression occurred in the lumen of tumor blood vessels (red arrows) in BC tissue from patients.

